# Zika virus dumbbell-1 structure is critical for sfRNA presence and cytopathic effect during infection

**DOI:** 10.1101/2023.01.23.525127

**Authors:** Monica E. Graham, Camille Merrick, Benjamin M. Akiyama, Matthew Szucs, Sarah Leach, Jeffery S. Kieft, J. David Beckham

## Abstract

Zika virus (ZIKV) contains multiple conserved RNA structures in the viral 3’ untranslated region (UTR), including the structure known as dumbbell-1 (DB-1). Previous research has shown that the DB-1 structure is important for flavivirus genome replication and cytopathic effect (CPE). However, the role of the DB structure and the mechanism by which it contributes to viral pathogenesis is not known. Using recently solved flavivirus DB RNA structural data, we designed two DB-1 mutant ZIKV infectious clones termed ZIKV-TL.PK, which disrupts DB-1 tertiary folding and ZIKV-p.2.5’, which alters DB-1 secondary structure formation. In cell culture, we found that viral genome replication of both mutant clones is not significantly affected compared to ZIKV-WT, but viral CPE is considerably decreased. We investigated sub-genomic flaviviral RNA (sfRNA) formation by both DB-1 mutants following A549 infection and found both mutant clones have decreased levels of all sfRNA species compared to ZIKV-WT during infection. To investigate the mechanism of decreased CPE in our DB-1 mutant clones, we assayed ZIKV DB mutant-infected A549 cells for cell viability and caspase activation. We found that cell viability is significantly increased in DB-1 mutant-infected cells compared to ZIKV-WT due to reduced caspase 3 activation. We also show that replication of the ZIKV-P.2.5’ mutant was significantly restricted by type I interferon treatment without altering interferon stimulated gene expression. Using a murine model of ZIKV infection, we show that both ZIKV-DB-1 mutants exhibit reduced morbidity and mortality compared to ZIKV-WT virus due to tissue specific attenuation in ZIKV-DB viral replication in the brain tissue. Overall, our data show that the flavivirus DB-1 RNA structure is important for maintaining sfRNA levels during infection which supports caspase-3 dependent, viral cytopathic effect, type 1 interferon resistance, and viral pathogenesis in a mouse model.

## Introduction

The *Flavivirus* genus contains multiple mosquito-borne RNA viruses that impact human health, including West Nile virus (WNV), Dengue virus (DENV), yellow fever virus (YFV), and Zika virus (ZIKV)^1^. ZIKV, an emerging global pathogen, was first discovered in 1947 in Uganda, and has since spread across Africa, southeastern Asia, Micronesia, and South America^2–9^. It was discovered during the 2015-2016 Brazil outbreak that ZIKV could be transmitted vertically mother-to-fetus and cause a spectrum of congenital abnormalities referred to as “congenital Zika syndrome”^10^. While ZIKV has caused significant outbreaks in more equatorial regions of the world, changing climate and human distribution could expand geographic range and outbreaks^11^. With no currently approved antivirals or vaccines for ZIKV, the virus still poses a significant risk to human health.

Flaviviruses, including ZIKV, contain in their 5’ and 3’ untranslated regions (UTRs) multiple conserved structural motifs that have evolved to support viral growth and pathogenicity (Figure 1)^12,13^. Conserved 5’UTR structures are critical for NS5 binding and activity as well as ribosome subunit positioning around the start codon^12^. The 3’UTR contains several RNA structures with a wide range of functions that include flaviviral noncoding RNA production, cis-acting RNA genome interactions that support genome replication, and RNA-protein interactions^12,13^. While there is slight variation between individual viruses, the overall RNA structural organization of the 3’UTR is consistent with one or more structures known as exonuclease-resistant RNAs (xrRNAs), two dumbbells (DB), short hairpin (sHP), and 3’ stem loop (3’SL)^12^. At the 5’ end of the 3’UTR, the xrRNAs are crucial for the formation of sub-genomic flaviviral RNAs (sfRNAs)^12,14,15^. At the 3’ end of the 3’UTR are the sHP and 3’SL, which play an important role in viral non-structural protein binding to the 3’ end of the genome^13^. Between the xrRNAs and 3’SL lie the DB structures. DENV and WNV contain two complete DB structures, while YFV and ZIKV contain one complete DB, and one incomplete DB, referred to in this publication as a pseudo-dumbbell^12,16^. Of the flavivirus 3’UTR RNA structures, the DBs are the least thoroughly studied, both in terms of structure-function relationships and function during infection. In this study, we sought to investigate the contribution of the DB-1 structure to ZIKV pathogenesis. We created two mutant ZIKV clones disrupting the integrity of secondary and tertiary DB-1 structural elements. We found that our disruptions had minimal effects on genome replication but had significant effects on reducing viral cytopathic effect *in vitro*. We found that decreased cytopathic effect was due to a marked reduction in sfRNA levels during DB-1 mutant infection. DB-1 mutant clones also exhibited increased type I interferon sensitivity, and decreased caspase 3 activation during infection. In a murine model of ZIKV infection, we found that DB-1 mutants exhibited a modest decrease in genome replication but a marked increase in survival of DB-1 mutant-infected mice compared to ZIKV-WT infected mice. Our findings show for the first time that the ZIKV DB-1 structural integrity is correlated with the presence of sfRNAs during infection, which in turn correlates with ZIKV cytopathic effect *in vitro* and pathogenesis *in vivo*.

**Figure 1:**
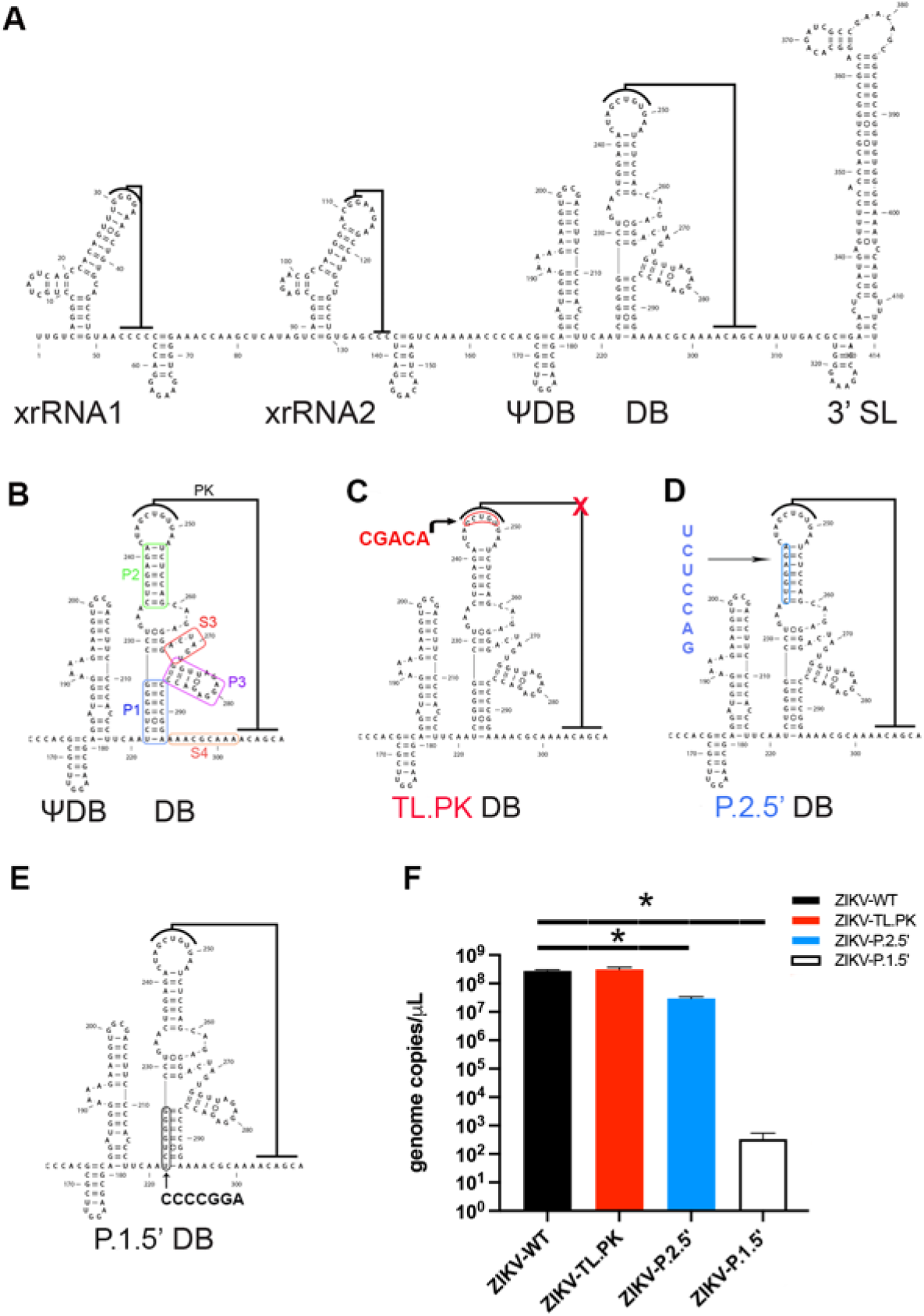
Zika virus Dumbell (DB) mutation design and clone-derived virus rescue. **A)** Schematic of the Zika virus (ZIKV) 3’ untranslated region (UTR) RNA secondary structural organization. **B)** Individual stem-loop designations in the DB structure. **C)** Mutational target in the DB structure to create the ZIKV-TL.PK mutant. Red=mutation target. **D)** Mutational target in the DB structure to create the ZIKV-P.2.5’ mutant. Blue=mutation target. **E)** Mutational target in the DB structure to create the ZIKV-P.1.5’ mutant. Black=mutation target. **F)** Genome copies of indicated wild-type (WT) or ZIKV mutant virus in the supernatant following transfection and virus rescue. *p<0.0004, ANOVA with Dunnett’s multiple comparisons test. N=3 per group.

## Results

### Generation and Rescue of ZIKV-TL.PK and ZIKV-p.2.5’ Infectious Clones

The ZIKV DB-1 structure is predicted to contain a pseudoknot fold, based on the 3D structure of the 3’UTR dumbbell of the insect-specific Donggang virus^17^. Specifically, the apical loop on the P2 stem forms Watson-Crick base pairs with downstream 3’UTR sequence (Figure 1A,B). In mosquito-borne flaviviruses this loop pairs with the 3’ cyclization sequence, which can pair with the upstream 5’ cyclization sequence to allow for genome replication^13^. Secondary structure analysis has also revealed that the ZIKV DB-1 structure very likely folds the same way as other flaviviral 3’UTR dumbbells^16,17^. For our first ZIKV mutant clone, termed TL.PK in this study, we mutated the pseudoknot-forming apical loop on the P2 stem (Figure 1B). We substituted this apical loop sequence with its Watson-Crick complement, eliminating the possibility of pseudoknot base pairing (Figure 1C). Our second and third ZIKV mutant clones, termed p.1.5’ and p.2.5’, followed the same rationale and design targeting the conserved stems in the DB-1 P1 and P2 stems, respectively (Figure 1D,E). mFold secondary RNA structure prediction software that both mutations result in significant disruptions to the DB-1 secondary structure (Supplementary Figure 1)^18^.

From the stock virus created for each ZIKV DB-1 mutant clone, we assessed viral genome copy number, infectious titer, and stability of mutations through Vero and C6/36 passaging. Quantifying genome copy numbers by RT-qPCR revealed that the ZIKV-TL.PK mutant exhibited similar genome copy numbers to ZIKV-WT (Figure 1F). The ZIKV-p.2.5’ clone exhibited a significant 9.3-fold reduction in genome copy numbers compared to ZIKV-WT, but still replicated to within a log10 of ZIKV-WT and ZIKV-TL.PK. However, the ZIKV-p.1.5’ exhibited mean decrease of 10^6^ genome copies compared to ZIKV-WT. Based on these results, we determined that, at least for viral stock generation, the ZIKV-TL.PK and ZIKV-p.2.5’ replicated sufficiently for downstream characterization. However, we determined that the p.1.5’ clone was unable to replicate sufficiently for evaluation in these studies. Following growth of stock ZIKV-WT, ZIKV-TL.PK and ZIKV-p.2.5’, we sequenced the 3’UTR and found unchanged mutations in the individual isolates. (Supplementary Figure 2).

### In Vitro Viral Kinetics of ZIKV-TL.PK and ZIKV-p.2.5’ Infectious Clones

To understand potential phenotypic effects of the TL.PK and p.2.5’ mutants, we analyzed the viral kinetics of our ZIKV-TL.PK and ZIKV-p.2.5’ clones in vitro. Mammalian A549 cells were infected at an MOI = 0.1 and samples were harvested at 0, 24, 48, and 72 hours post infection (hpi). We analyzed production of new infectious virus in the supernatant, as well as (+)-strand genome replication in supernatant and cell-associated samples (Figure 2). In both supernatant-associated and cell-associated samples, we observed up to an average ten-fold decrease in ZIKV-TL.PK and ZIKV-P.2.5’ genome replication at 24 hpi compared to ZIKV-WT (Figure 2A,B), but these values were determined not to be significantly different from ZIKV-WT genome replication. Additionally, we found no significant difference in quantity of genome replication between our two mutants and ZIKV-WT at 48 and 72 hpi.

**Figure 2:**
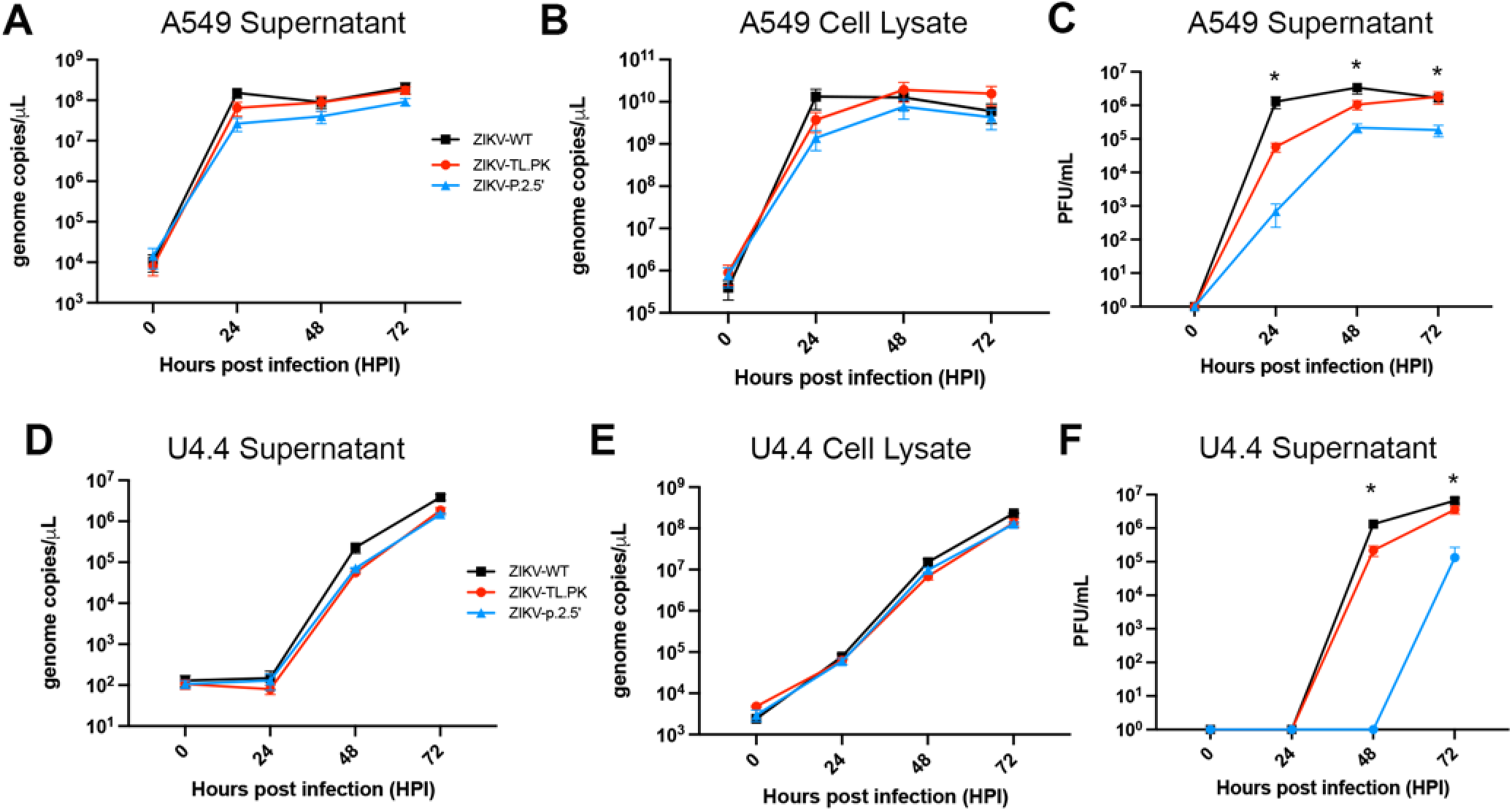
ZIKV DB mutants exhibit moderately reduced infectious virus titer in cell culture. A549 cells were inoculated with ZIKV-WT, ZIKV-TL.PK, and ZIKV-P.2.5’ (MOI 0.1) followed by harvest of **A)** supernatant and **B)** A549 cells at the indicated time points post-infection. C) A549 cells were inoculated with ZIKV-WT, ZIKV-TL.PK, and ZIKV-P.2.5’ (MOI 0.1) followed by harvest of supernatant for plaque assay analysis. *p<0.0001, Two-way ANOVA. N=9 total per group/time point, completed as triplicates in 3 experimental replicates. U4.4 cells were inoculated with ZIKV-WT, ZIKV-TL.PK, and ZIKV-P.2.5’ (MOI 0.1) followed by harvest of **D)** supernatant and **E)** U4.4 cells at the indicated time points post-infection. **F)** U4.4 cells were inoculated with ZIKV-WT, ZIKV-TL.PK, and ZIKV-P.2.5’ (MOI 0.1) followed by harvest of supernatant for plaque assay analysis. *p<0.0001, Two-way ANOVA. N=3 per group/time point.

Supernatant from all samples was also analyzed by plaque forming unit (PFU) assay to measure production of new infectious virus during infection. At 24 hpi, ZIKV-TL.PK and ZIKV-P.2.5’ produced over 10-fold and 1000-fold, respectively, less infectious virus compared to ZIKV-WT at the same time point (Figure 2C). ZIKV-TL.PK virus production caught up to ZIKV-WT by 72 hpi, while ZIKV-P.2.5’ consistently produced 10-fold less infectious virus than ZIKV-WT and ZIKV-TL.PK at later time points.

To understand the phenotypic effects of the TL.PK and p.2.5’ mutants in invertebrate cells, we infected U4.4 mosquito cells with ZIKV-WT, ZIKV-TL.PK and ZIKV-p.2.5’ following the same conditions stated above. We found that ZIKV-TL.PK and ZIKV-p.2.5’ exhibited unchanged genome copy replication from ZIKV-WT in the supernatant and inside cells at all time points (Figure 2D,E). As in mammalian cells, we found that production of infectious ZIKV was similar in U4.4 cells for ZIKV-WT and ZIKV-TL.PK isolates, and the ZIKV-TL.PK mutant catches up in infectious virus production at 72 hours (Figure 2F). ZIKV-p.2.5’ exhibited significant reduction in infectious virus titer production that was undetectable through 48 hours but increased at 72 hours while still exhibiting a 50-fold reduction at 72 hours compared to ZIKV-WT (Figure 2F).

In our evaluation of viral titer in the ZIKV mutants, we noted that the plaque morphology of ZIKV-TL.PK and ZIKV-P.2.5’ was altered due to reduced cell monolayer clearance, resulting in less well-defined plaque edges compared to ZIKV-WT (Figure 3A). These data align with findings from previous studies that showed flaviviruses with 3’UTR structural mutations produced reduced cytopathic effect and plaque sizes compared to WT virus^14,19–21^. We next wanted to quantify the cytopathic effect of ZIKV-TL.PK and ZIKV-p.2.5’ clones in A549 cells using an XTT cell viability assay. A549 cells were inoculated with ZIKV-WT, ZIKV-TL.PK and ZIKV-p.2.5’ (MOI = 0.1) and harvested at 24, 48, and 72 hpi for analysis. ZIKV-WT infected cells exhibited a significant decrease in cell viability throughout all time points compared to mock infected control cells (Figure 3B). Cells infected with ZIKV-TL.PK exhibited an initial 24% reduction in viability compared to mock-infected control cells at 24 and 48 hpi. However, ZIKV-TL.PK infected cells exhibited no significant difference in cell viability at 72 hpi compared to controls cells (Figure 3B). Cells infected with ZIKV-p.2.5’ mutants exhibited no significant change in cell viability compared to mock-infected cells throughout the time course. (Figure 3B).

**Figure 3:**
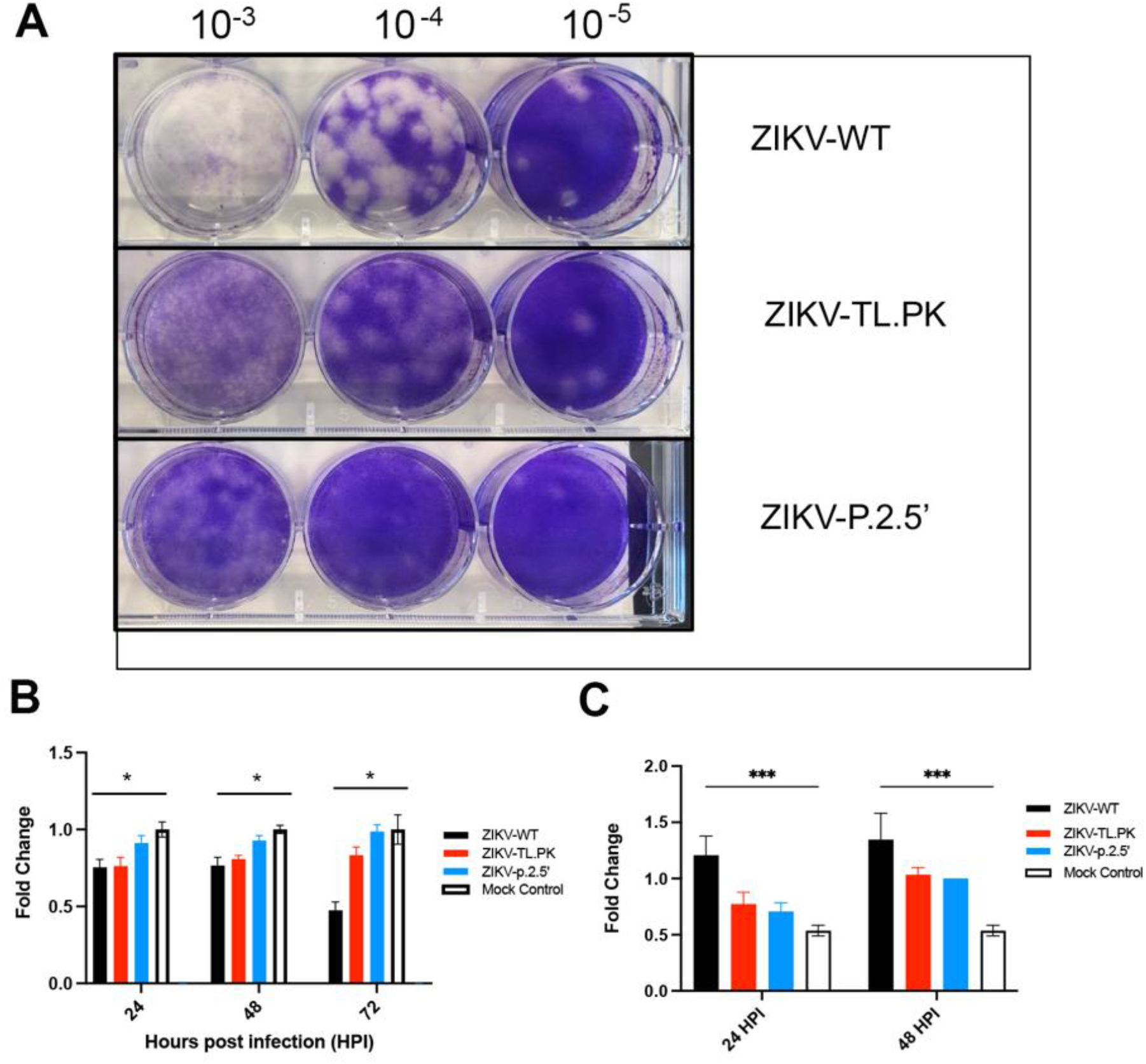
ZIKV-TL.PK and ZIKV-P.2.5’ exhibit reduced pathogenesis due to reduced caspase-3 activation. **A)** Image of representative plaque assays for ZIKV-WT, ZIKV-TL.PK, and ZIKV-P.2.5’ showing altered plaque phenotype in ZIKV DB mutants. A549 cells were inoculated with ZIKV-WT, ZIKV-TL.PK, and ZIKV-P.2.5’ (MOI 0.1) and harvested at indicated time points for **B)** XTT assay (N=9 per group/time point) and **B)** Caspase-3 activity assay (N=3 per group/time point). 2way ANOVA, *p=0.0001, ***p<0.0001.

To define the mechanism by which ZIKV-TL.PK and ZIKV-p.2.5’ cause reduced cell injury, we inoculated A549 cells with ZIKV-WT, ZIKV-TL.PK and ZIKV-p.2.5’ (MOI 0.1) and evaluated caspase-3 and caspase-1 activation. Compared to ZIKV-WT inoculated cells, we found that ZIKV-TL.PK and ZIKV-p.2.5’ exhibited significantly less caspase 3 activation (Figure 3C). However, there was no significant change in caspase-1 activation following ZIKV infection in A549 cells (Supplementary Figure 3). These data show that ZIKV-TL.PK and ZIKV-p.2.5’ are attenuated due to decreased caspase-3 activation following infection.

### sfRNA Analysis of ZIKV-TL.PK and ZIKV-p.2.5’ Infectious Clones

Sub-genomic flaviviral RNA (sfRNA) formation and function are essential to flaviviral cytopathic effect and pathogenesis^15^. In the ZIKV 3’UTR, stem loops 1 and 2, also known as xrRNAs, are known to be crucial for the formation of sfRNAs^14^. There are also studies indicating that the flavivirus DB structures affect sfRNA levels in WNV, DENV, and ZIKV^17,19,20^. We infected A549 cells with ZIKV-TL.PK and ZIKV-p.2.5’ isolates (MOI = 0.25) and harvested cellular RNA at 12, 24, and 48 hpi to assay sfRNA formation using Northern blot analysis with a probe specific for the conserved region of the ZIKV DB-1 structure. In cells infected with ZIKV-WT, sfRNA bands start appearing at 12 hpi, with bands for sfRNAs 1, 2, and 3 at 24 and 48 hpi (Figure 4A). However, ZIKV-TL.PK mutant exhibited reduced levels of all sfRNAs at 24 and 48 hpi compared to WT. ZIKV-p.2.5’ exhibited no visible sfRNA production at 12 or 24 hpi, and faint bands for sfRNAs 1 and 2 at 48 hpi (Figure 4B). We also found no evidence of sfRNA3 production in either ZIKV TL.PK or ZIKV p.2.5’. Densitometry analysis using the ratio of sfRNA density corrected for genomic RNA density shows that there is a significant reduction of sfRNA levels in from both ZIKV-TL.PK and ZIKV-p.2.5’ compared to ZIKV-WT virus (Figure 4C). These data show that mutation of DB-1 3-D RNA structure results in marked decrease of sfRNA levels during infection independent of genome replication.

**Figure 4:**
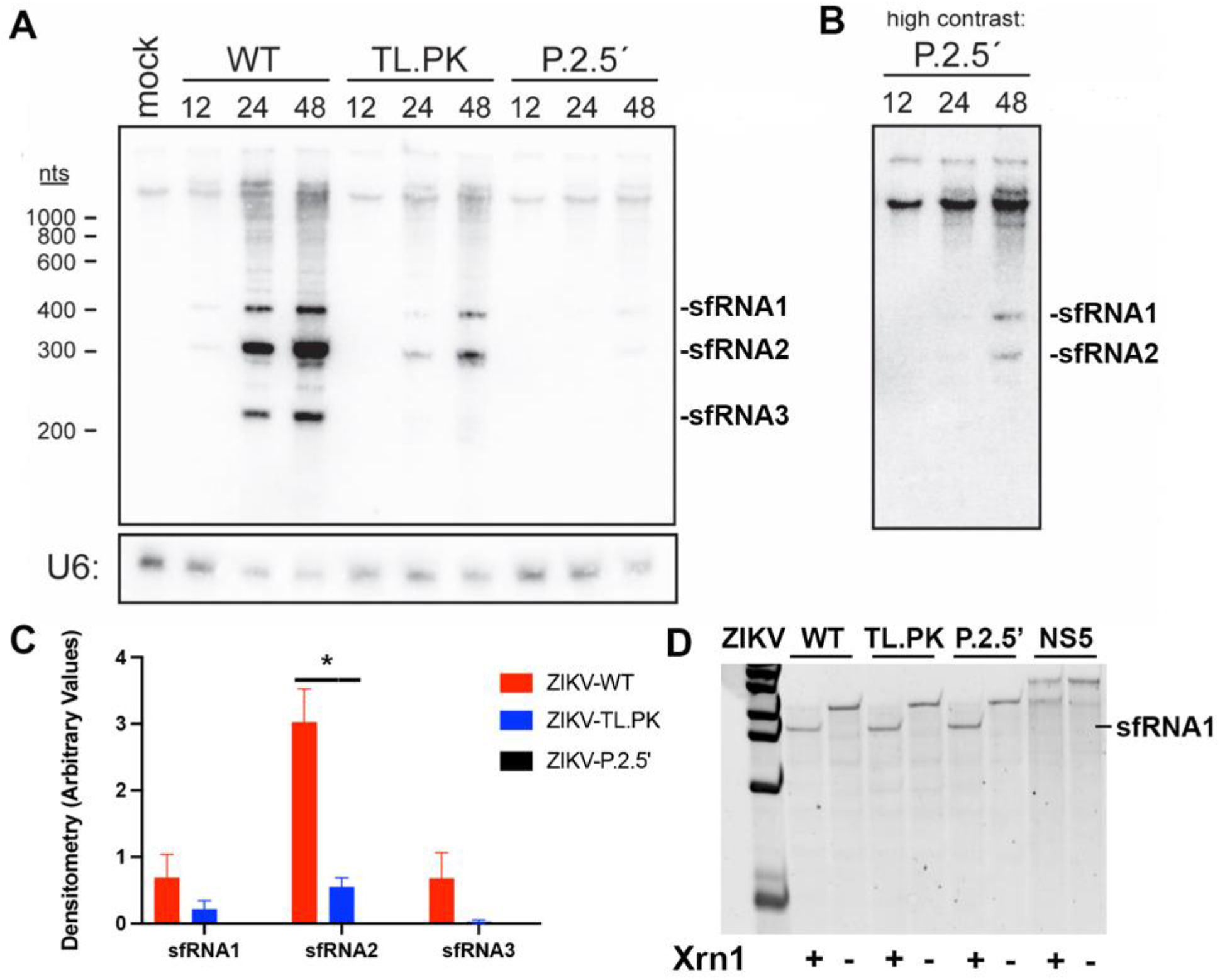
ZIKV DB mutations result in altered sfRNA expression. A549 cells were inoculated with ZIKV-WT, ZIKV-TL.PK, and ZIKV-P.2.5’ (MOI 0.1) and cells were harvested at indicated time points for **A)** northern blot analysis of sfRNA biogenesis at indicated hours post-infection. **B)** Enhanced image of ZIKV-P.2.5’ mutant northern blot image. **C)** Densitometry analysis of 24 hour time point of Northern blot bands. N=2 per group. *p<0.0001. D) Xrn-1 digest assay of 3’UTR from ZIKV-WT, ZIKV-TL.PK, ZIKV-P.2.5’, and RNA from ZIKV NS5 gene as a control. Representative image of N=2 per group.

To investigate whether ZIKV-TL.PK and ZIKV-p.2.5’ cause a loss of Xrn-1 resistance during infection, we performed an *in vitro* Xrn-1 digest of 3’UTR sequences containing either the WT, TL.PK, or p.2.5’ DB-1 sequence. We found no significant difference in production of sfRNA following Xrn-1 digest of 3’UTR sequences from ZIKV-TL.PK, and ZIKV-p.2.5’ compared to ZIKV-WT (Figure 4D). These data suggest that biogenesis of xrRNA1&2-dependent sfRNA1&2 is unchanged in ZIKV-TL.PK and ZIKV-p.2.5’, but overall sfRNA levels are decreased during infection following production.

### Mechanism of Reduced Cytopathic Effect In Vitro

sfRNAs have a myriad of functions during flavivirus infection, including contributing to viral cytopathic effect. Based on the sfRNA data for our ZIKV-TL.PK and ZIKV-P.2.5’ clones, we wanted to investigate which sfRNA functions were associated with the observed loss of cytopathic effect and caspase-3 activation.

An important function of sfRNAs during infection is to antagonize the type I interferon (T1I) responses to promote viral immune escape and viral replication^22–25^. It has been shown in previous studies that flaviviruses lacking sfRNAs are more susceptible to clearance by T1I during infection^21,26^. Since ZIKV-TL.PK and ZIKV-p.2.5’ exhibited reduced sfRNA expression compared to ZIKV-WT, we next evaluated sensitivity of ZIKV growth to T1I treatment. We infected Vero cells with ZIKV-WT, ZIKV-TL.PK, and ZIKV-p.2.5’ (MOI 0.1) followed by treatment with a dilution series of IFN-β and harvested cells at 48 hpi for viral genome quantification by RT-qPCR. We found that ZIKV-TL.PK (IC50=99.6 IU/ml) was not significantly more sensitive to interferon compared to ZIKV-WT (IC50=96.1 IU/ml). We did observe that ZIKV-p.2.5’ (IC50=50.07 IU/ml) was significantly more sensitive to interferon treatment compared to ZIKV-WT (Figure 5A, Supplementaary Figure 4).

**Figure 5:**
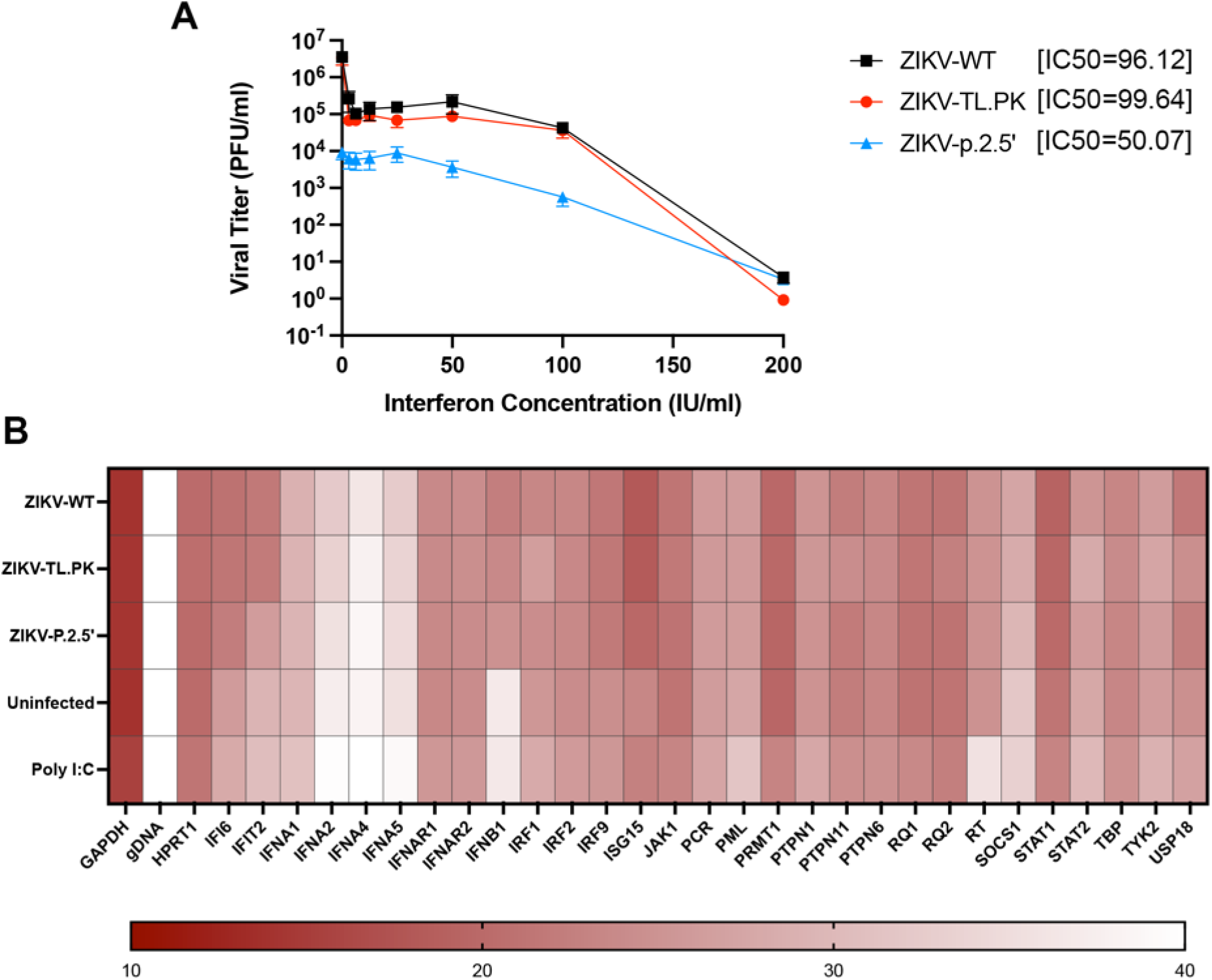
ZIKV DB mutant viruses exhibit increased sensitivity to interferon independent of interferon stimulated gene expression. **A)** Vero cells were inoculated with ZIKV-WT, ZIKV-TL.PK, and ZIKV-P.2.5’ (MOI 0.1) followed by treatment with type I interferon (a2) at indicated concentrations and harvested at 48 hours post-infection for plaque assay analysis. N=6 per virus/interferon treatment group. Calculated type 1 interferon IC50 for ZIKV-WT (96.12), ZIKV-TL.PK (99.64), and ZIKV-P.2.5’ (50.07). **B)** A549 cells were inoculated with ZIKV-WT, ZIKV-TL.PK, and ZIKV-P.2.5’ (MOI 0.1) and harvested at 24 hours post-infection for RT-PCR array for interferon-stimulated gene expression (N=6 per group).

The presence of sfRNAs can also significantly influence the induction of interferon stimulated genes (ISGs) during infection. sfRNA-deficient ZIKV has been shown to increase mRNA expression of several ISGs in the T1I pathway during in vitro infection, indicating that sfRNAs can play a crucial role in antagonizing the interferon response during infection^27^. We chose to investigate how ISGs in the interferon α/β pathway would be affected by the reduction of sfRNAs found in ZIKV-TL.PK or ZIKV-p.2.5’ mutants. A549 cells were inoculated with ZIKV-WT, ZIKV-TL.PK or ZIKV-p.2.5’ (MOI=0.01) and harvested 24 hpi for total RNA. Total RNA was then assayed by RT-qPCR for relative interferon α/β pathway ISG mRNA levels. At 24 hours post-infection, ISG mRNA levels were not significantly changed between cells infected with ZIKV-WT versus ZIKV TL.PK and ZIKV p.2.5’ mutants (Figure 5B). In ZIKV-p.2.5’ infected cells, mRNA levels for several ISGs were reduced compared to ZIKV-WT infected cells. We determined that, at 24 hpi, the absence of WT sfRNA levels in ZIKV-TL.PK and ZIKV-p.2.5’ does not significantly correlate to alterations in ISG mRNA expression compared to ZIKV-WT infection (Supplementary Table 1).

### Analysis of ZIKV-TL.PK and ZIKV-p.2.5’ in Murine Model

Studies have shown that deleterious mutations, as well as RNA structure-altering mutations in the flavivirus 3’UTR led to reduced viral burden and pathogenesis in mouse models^14,21,26,28–31^. To investigate potential alterations in the pathogenesis of our ZIKV DB-1 mutant clones, we used an AG129 mouse model for ZIKV infection. One cohort of mice were infected with 10^4^ PFU via intraperitoneal injection with ZIKV-WT, ZIKV-TL.PK, ZIKV-P.2.5’, or PBS and followed post-infection for weight loss and development of neuroinvasive disease. Mice were sacrificed when moribund over a 16-day time course. We found that ZIKV-TL.PK and ZIKV-P.2.5’ infected mice experienced significantly decreased weight loss compared to ZIKV-WT infected mice (Figure 6A, Supplementary Figure 5). Survival curve analysis showed that ZIKV-TL.PK (median survival=14.5 days) and ZIKV-P.2.5’ (median survival=15.5 days) infected mice exhibited a significantly longer survival time compared to ZIKV-WT (median survival = 9 days) infected mice (Figure 6B).

**Figure 6:**
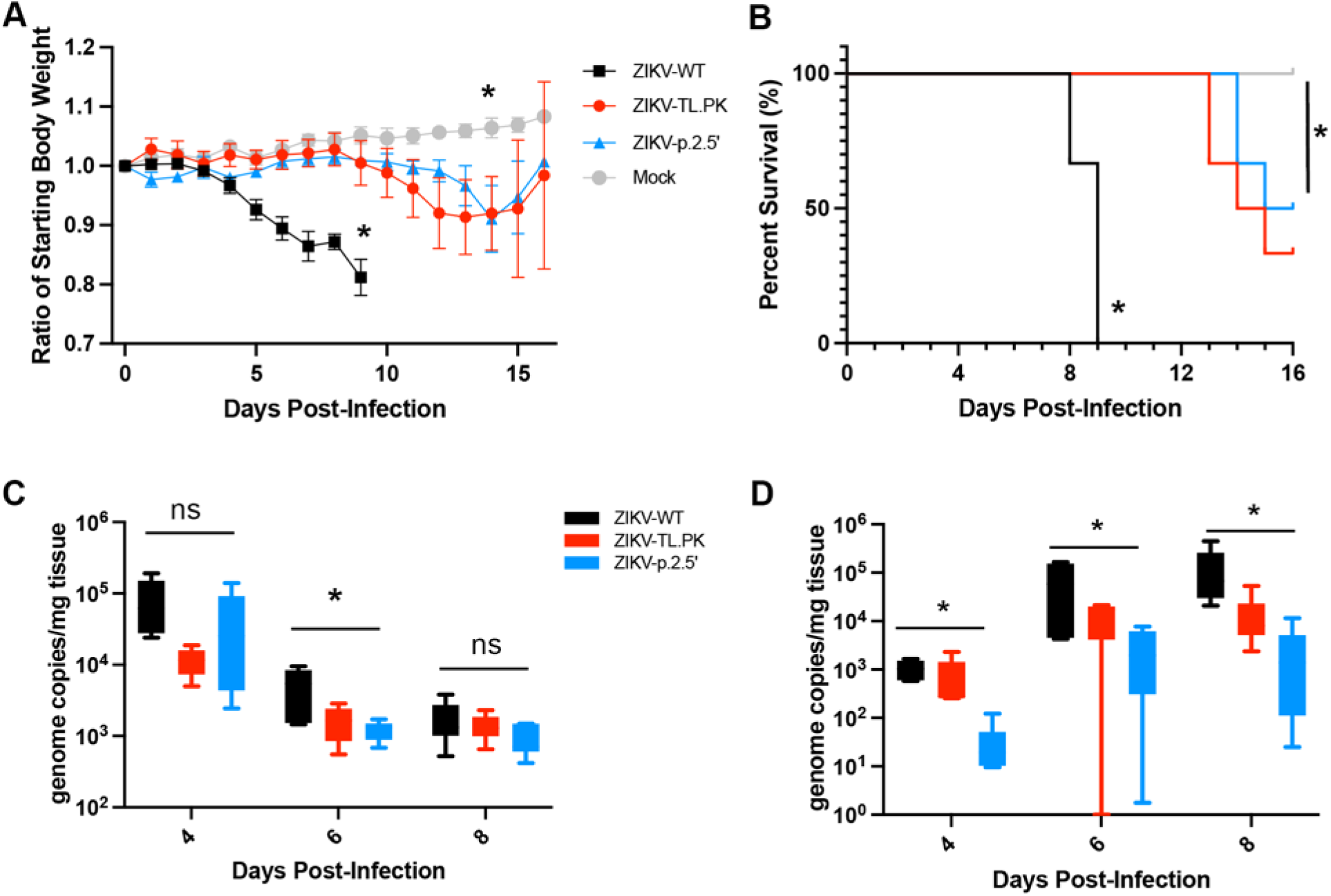
ZIKV DB mutants exhibit marked attenuation in AG129 mice. Mice were inoculated with ZIKV-WT, ZIKV-TL.PK, and ZIKV-P.2.5’ at 1000 pfu by intraperitoneal injection. **A)** Daily weights were recorded over 16 days (*p=0.0023, mixed effects analysis), and **B)** mice were euthanized when moribund (*p<0.0001, Log-rank test). N=6 per group. Mice were inoculated with ZIKV-WT, ZIKV-TL.PK, and ZIKV-P.2.5’ at 1000 pfu by intraperitoneal injection followed by tissue harvest at the indicated time point post-infection (N=6 per group per time point). ZIKV genome copies were determined in the **C)** spleen (*p=0.0115) and **D)** brain (*p<0.048), One-way ANOVA.

Next, AG129 mice were infected as described above and sacrificed 4-, 6-, or 8-days post-infection to analyze ZIKV genome copies in the spleen and brain. We found no significant difference in genome replication in the spleen when comparing ZIKV-WT to ZIKV-TL.PK or ZIKV-p.2.5 genome replication throughout the time course (Figure 6C). However, we found a significant reduction (p=0.03) in ZIKV-TL.PK and ZIKV-p.2.5 genome replication in the brain compared to ZIKV-WT inoculated mice throughout the time course of infection (Figure 6D). These data show that the ZIKV-TL.PK and ZIKV-p.2.5’ clones exhibit marked, end-organ specific attenuation in an immune compromised murine model of ZIKV infection.

## Discussion

In this study, we sought to understand the contribution of the ZIKV 3’UTR DB-1 to viral pathogenesis. We utilized the recently resolved DB 3-D structure from Donggang virus to guide targeted mutations to disrupt the predicted secondary and tertiary structure of DB-1^17^, with the goal of eliminating the native role of DB-1 structure while maintaining the full-length sequence of the 3’UTR. Based on previous studies and hypothesized functions of flaviviral dumbbell structures, we initially hypothesized that the ZIKV DB-1 structure was important for regulating viral genome replication, and that disrupting the structural integrity would alter viral genome replication. When analyzing viral kinetics of ZIKV-TL.PK and ZIKV-P.2.5’ in cell culture, we found that genome replication was not significantly altered compared to ZIKV-WT virus. However, we did find evidence of decreased infectious virion production, as well as significant attenuation *in vitro* and *in vivo*. We also show that attenuation of ZIKV-TL.PK and ZIKV-P.2.5’ is closely associated with decreased sfRNA levels, which correlates with decreased caspase-3 activation and increased type 1 interferon sensitivity. These data show for the first time that the structural integrity of DB-1 is important to support sfRNA levels following infection, as well as pathogenic effects associated with sfRNAs.

Sub-genomic flaviviral RNAs (sfRNAs) are non-coding RNA made from the viral 3’UTR after resisting host Xrn-1. The role of xrRNA1 and xrRNA2 are the main structures responsible for 3’UTR Xrn-1 resistance, and their complex tertiary folding is critical for their ability to resist Xrn-1 degradation resulting in biogenesis of sfRNA1 and sfRNA2, respectively^14,32,33^. Previous work to resolve the structure of the Donggang virus dumbbell found that while the DB forms a pseudoknot, it does not make the same loop around the 5’ end of the RNA to confer Xrn-1 resistance like xrRNA1 and xrRNA2.^17^ Prior studies show that the ZIKV DB-1 does not efficiently resist Xrn-1 degradation on its own, and that sfRNA3 is likely generated by 3’ end processing and not DB-1 resisting Xrn-1^17^. In this study, the ZIKV-TL.PK and ZIKV-P.2.5’ clones exhibited a significant decrease in levels of all sfRNA species compared to ZIKV-WT, and both clones exhibited loss of sfRNA3 levels. We also found that *in vitro* Xrn-1 digestion of ZIKV-TL.PK and ZIKV-P.2.5’ 3’UTR sequences exhibited no changes in sfRNA production. Our findings suggest that DB-1 plays a neutral role in Xrn-1-mediated sfRNA biogenesis such that structural changes that disrupt the 3-D structure of DB-1 do not alter sfRNA formation. Based on our and prior findings, we hypothesize that the DB-1 structure prevents sfRNAs from undergoing additional 3’ exonuclease or endonuclease degradation, and loss of the DB-1 structural integrity results in increased sfRNA turnover.

sfRNAs have a myriad of functions during infection, and sfRNA-deficient flaviviruses display decreased cytopathic effect and pathogenesis. We hypothesized the loss of cytopathic effect following infection with ZIKV-TL.PK and ZIKV-P.2.5’ clones was due to decreased sfRNA levels, since there were otherwise no significant changes in genome replication. Studies using infection flavivirus clones with or without sfRNAs showed reduced activation of apoptosis, specifically caspase-3, during in vitro infection^20,34^. A previous study using DENV2 specifically showed that reconstitution of WT sfRNA levels partially recovered caspase-3 activation back to levels seen with DENV2-WT^20^. Similarly, we found that ZIKV-TL.PK and ZIKV-P.2.5’ clones that expressed an overall reduction in sfRNA production following infection exhibited significantly decreased caspase-3 activation, but not caspase-1 activation, compared to ZIKV-WT. Our results support an important role for flavivirus sfRNA levels in caspase activation as a mechanism of virus-induced cytopathic effect.

Our studies with the ZIKV-P.2.5’ clones show that alteration of the 3-D DB-1 structure and decrease in sfRNA levels results in increased sensitivity to (T1I) treatment compared to ZIKV-WT. Using clone-derived deletion models of the DBs, previous studies have shown similar effects of T1I on sfRNA-deficient flaviviruses. WNV and YFV lacking sfRNAs replicated better in cells lacking factors in the T1I response than cells competent for all T1I factors^19,26^. A prior study also demonstrated that exogenous addition of T1I had a significant negative effect on virus replication, suggesting that sfRNA-deficient viruses are more sensitive to T1I^26^. Our data showed that the ZIKV-TL.PK did not exhibit significantly altered IFN sensitivity compared to ZIKV-WT, however ZIKV-P.2.5’ exhibited significantly increased IFN sensitivity. While both clones exhibit significantly less sfRNA compared to ZIKV-WT, the ZIKV-P.2.5’ exhibited a greater reduction in sfRNA levels. It may be that even low levels of sfRNA during ZIKV-TL.PK infection was able to antagonize T1I responses, whereas ZIKV-P.2.5’ sfRNA levels are below a threshold needed to antagonize the interferon response. More studies comparing the sfRNA expression levels with interferon signaling are needed to define a potential dose-effect for sfRNA production to alter T1I signaling.

In our studies of ISGs in the IFN α/β pathway, we found that ISGs are upregulated following ZIKV infection but there were no significant changes in expression exhibited between ZIKV WT, ZIKV-TL.PK and ZIKV-P.2.5’ infected cells. These data suggest that sfRNA production does not alter individual ISG mRNA expression. The increased sensitivity of ZIKV-P.2.5’ may be due to individual interactions between sfRNAs and ISGs proteins directly, or sfRNAs and viral nonstructural proteins like NS5 are required to inhibit the T1I response during infection^34,35^. Further studies are needed to evaluate interactions between sfRNAs, viral proteins, and individual ISGs known to restrict effector functions of important flavivirus restriction factors.

To determine the attenuation of the ZIKV-TL.PK and ZIKV-P.2.5’, we inoculated AG129 mice and determined survival and viral genome copies post-infection. These data show marked attenuation of the ZIKV-TL.PK and ZIKV-P.2.5’ as evident by significantly reduced weight loss and prolonged survival following infection. While ZIKV-TL.PK and ZIKV-P.2.5’ clones replicated in the spleen to equivalent levels as ZIKV-WT, viral genome copies of ZIKV-TL.PK and ZIKV-P.2.5’ clones were significantly decreased in the brain. These data imply that mutation of the DB-1 structure and subsequent decrease in sfRNA levels result in decreased replication in the brain as a mechanism for the decreased mortality seen in this murine model. Future studies will need to determine virus-specific neutralizing antibody response and T-cell responses induced by ZIKV-TL.PK and ZIKV-P.2.5’ in immune competent models to determine the role of these attenuation approaches in future vaccine development. Further studies to determine attenuation and immunogenicity of DB mutants may provide a common attenuation approach for vaccine development against mosquito-borne flaviviruses by disrupting the DB-1 structure in the 3’UTR.

## Materials and Methods

### Cell Lines and Viruses

Mammalian Vero cells, mammalian A549 cells, and mosquito C6/36 cells were sourced from ATCC. Mosquito U4.4 cells were generously provided by Dr. Aaron Brault (CDC Center for Vector-Borne Diseases). Vero and C6/36 cells were cultured in 1X Minimal Essential Media (ThermoFisher), and A549 cells were cultured in Ham’s F-12 Nutrient Mix (ThermoFisher). Media for Vero, C6/36, and A549 cells were supplemented with 10 % FBS, 1X MEM non-essential amino acids (ThermoFisher), 100 μM sodium pyruvate (ThermoFisher), 1 mM HEPES (ThermoFisher), and 1X penicillin/streptomycin (company). U4.4 cells were cultured in Mistuhashi and Maramorosch (M&M) insect media, and supplemented with 10% FBS, 1X MEM non-essential amino acids, and 1X penicillin/streptomycin. Mammalian cell lines were cultured at 37°C with 5% CO_2_. Mosquito cell lines were cultures at 28°C with 5% CO_2_. Viruses used in this study are a wild-type ZIKV clone derived from the PRVABC59 ZIKV isolate, as well as two clone-derived ZIKV mutant strains^36^.

### Plasmids and Generation of TL.PK and p.2.5’ mutants

Previously described pACYC177 vector plasmids containing ZIKV PRVABC59 genome from either the 5’ UTR to nt 3498 (pJW231) or from nt 3109 to the end of the 3’ UTR (pJW232) were used to generate WT, TL.PK, and p.2.5’ ZIKV clones^36^. TL.PK and p.2.5’ mutations were cloned into the pJW232 plasmid with gBlocks (IDT) using Gibson assembly. Mutant gBlock inserts and pJW232 vector were linearized and amplified using PCR (Table 1). Gibson assembly was performed with the NEB Gibson Assembly® Master Mix (New England Biolabs) with a vector-to-insert ratio of 1:5. Assembled plasmids were transformed into and isolated from NEB Stable Competent *E. Coli* cells (New England Biolabs) and amplified with rolling cycle amplification. Mutations were verified with Sanger sequencing (Eton Biosciences, San Diego, CA, USA).

**Table 1:**
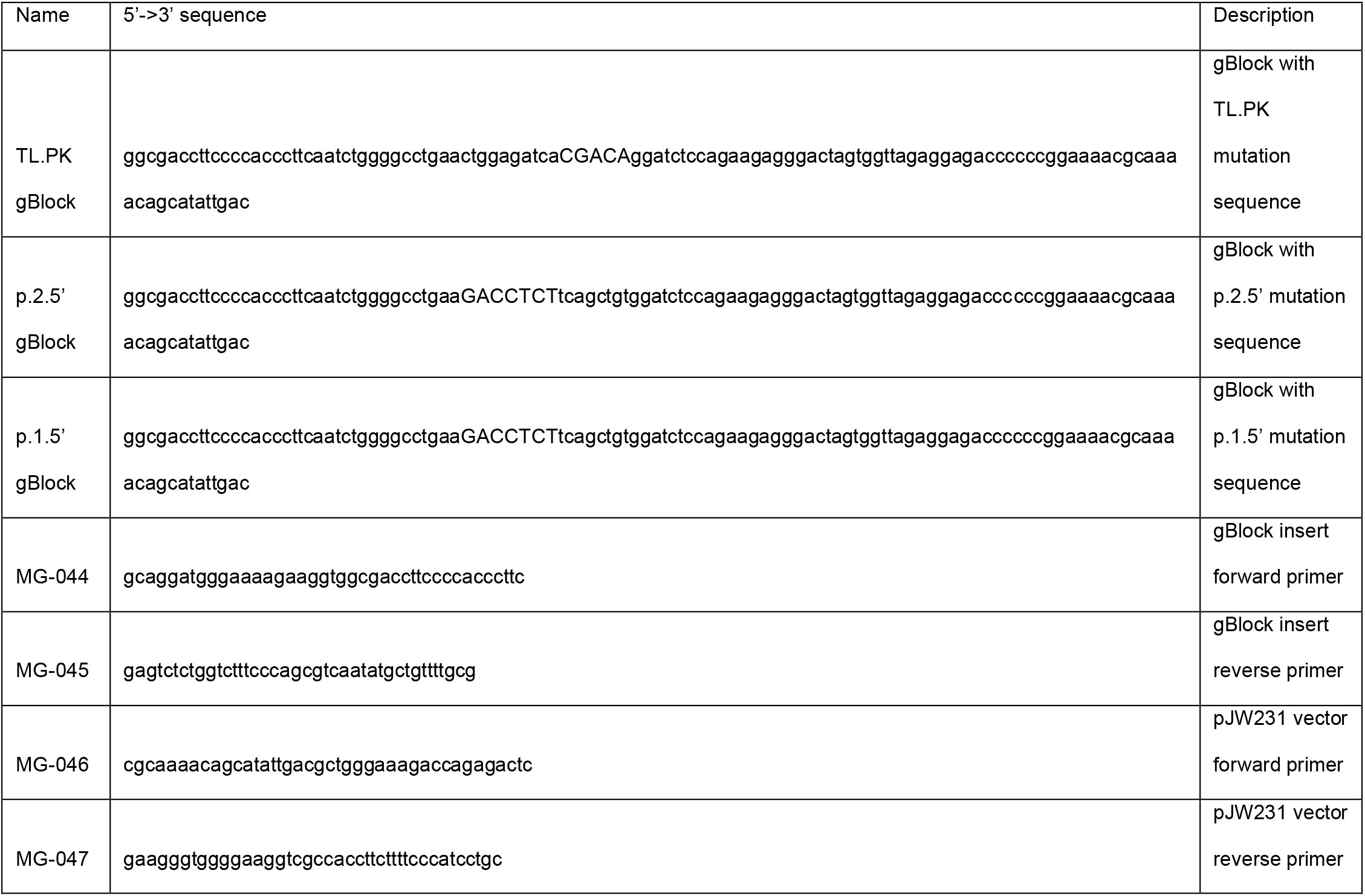
gBlock and primer sequences used for generating mutants.

### Propagation and Rescue of Zika Virus Clones

Wild type, TL.PK, and p.2.5’ pJW232 plasmids and wild type pJW231 plasmid were digested and ligated together to create a DNA template of the complete ZIKV genome according to Sparks et al^28^. Wild type, TL.PK, and p.2.5’ genomic RNA was generated by in vitro transcription using the HiScribe™T7 ARCA mRNA kit (New England Biolabs). For ZIKV-WT, genomic RNA was electroporated into mosquito C6/36 cells using the following protocol: 8e^6^ cells were suspended in 400 μL of PBS with 40 μL of in vitro transcribed RNA. Cells were electroporated in a 0.2 cm cuvette with a square wave with 3 μF capacitance and 750 V for a 1msec pulse. Cells were pulsed twice with five seconds between pulses. After electroporation cells were rested at room temperature for 15 minutes.

For the DB-1 mutants, 40 μg of TL.PK and p.2.5’ genomic RNA was transfected into Vero cells using MessengerMAX lipofectamine reagent (Invitrogen). Virus was harvested when approximately 70% cell clearance was observed. Supernatant was spun down to clarify, and FBS and HEPES was added to supernatant to final concentrations of 20% and 10 mM, respectively. Virus was aliquoted and stored at −80°C. Extracellular viral RNA was quantified as described below in “Virus Quantification”. TL.PK and p.2.5’ Vero stocks were then blind passaged in C6/36 cells to generated higher titer virus. C6/36 cells were plated in T-182 flasks to sub-confluency. Volume of virus with a calculated value of 1e^8^ viral genomes were added to each flask. Virus was harvested when approximately 70% cell clearance was observed. Virus was harvested and stored as previously described. Viral titer was determined via plaque forming unit assay as described in “Virus Quantification”.

### Virus Quantification

The cell-free infections virus from the supernatant of infected cells were quantified using plaque forming unit assay. Stock virus samples were serially diluted ten-fold 10^0^ to 10^-5^. 500 μL of each dilution was added to Vero cells plated in 6-well tissue culture plates to inoculated cells for one hour at 37°C. After inoculation cells and viral inoculum were overlayed with a 1:1 mixture of 2.5% Avicel in DI-H_2_O and 2X MEM supplemented with 10% FBS, 10 X sodium pyruvate, 10X non-essential amino acids, 10X penicillin-streptomycin, and 100 mM HEPES. Plaque assay plates were incubated for 6 days at 37°C. After incubation cells were washed with 1 mL 1X PBS, then fixed with 4% paraformaldehyde for 15 minutes at room temperature. After PFA removal plates were stained with 1% Crystal Violet in ethanol for one minute. Plates were washed three times with DI-H_2_O. Plaques were counted and stock virus titer (PFU/mL) was calculated using the following equation:

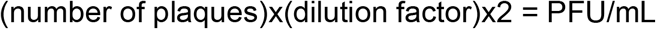

Extracellular viral RNA was isolated from infected cell supernatant using the E.Z.N.A Viral RNA kit (Omega-BioTek). Cell-associated viral RNA was extracted from infected cells using the E.Z.N.A Total RNA kit (Omega-BioTek). Viral RNA was reversed transcribed and quantified by qPCR.

### 3’UTR Sequencing

Viral genomic RNA for ZIKV-WT, TL.PK, and p.2.5’ was isolated from viral stocks using the OmegaBioTek E.Z.N.A Viral RNA Kit. Using the linker sequence and reverse transcription primer described in Table 2. Viral cDNA was generated following the linker pre-adenylation, linker ligation, and reverse transcription protocols from McGlincy 2017 (find to cite correctly). 3’UTR fragments were amplified using NEB Phusion Polymerase kit with the following parameters: 98°C for 30 seconds; 30 cycles of: 98°C for 10 seconds, 58°C for 15 seconds, 72°C for 15 seconds; 72°C for 10 minutes. Sanger sequencing was done by Eton BioSciences (San Diego, CA).

**Table 2:**
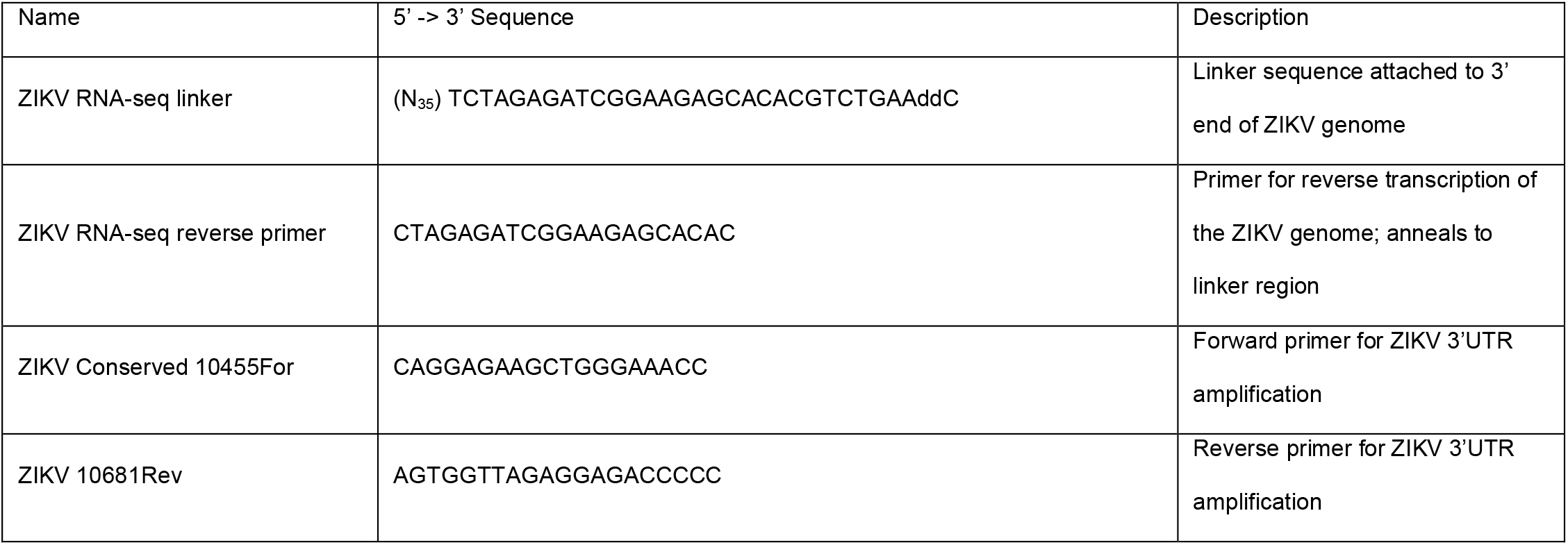
Linker and Primer Sequences for 3’UTR Sequencing.

### In Vitro Viral Growth Kinetics

Mammalian A549 or mosquito U4.4 cells were seeded in 6-well plates at a density of 2e^5^ and 6e^4^ cells per well, respectively. Cells were infected with WT, TL.PK, or p.2.5’ virus at an MOI = 0.1. At 0, 24, 48, and 72 hours post-infection, supernatant was harvested for quantification of viral genome copies by RT-qPCR, and production of infectious virus by focus forming unit assay. Infected cells were also harvested for quantification of cell-associated viral genome copies by RT-qPCR.

### XTT Assay

A549 cells were plated in a 96-well clear bottom cell culture plate at a density of 10^4^ cells per well. Cells were infected with ZIKV-WT, TL.PK, and p.2.5’ virus at an MOI of 0.1 for one hour at 37°C. Inoculum was remove and 200 μL of complete Ham’s F-12 media was added to each well. XTT assay was performed with the XTT Cell Viability Kit from Cell Signaling Technology. At the harvest time point, control wells for 0% viability were treated with 100 μL 4% PFA for 15 minutes. Media in all wells was adjusted to 100 μL of complete Ham’s F-12 media. 50 μL of XTT detection solution was added to each well and incubated at 37°C for four hours. After incubation, culture media in each well was treated with 100 μL 4% PFA for 15 minutes at room temperature to inactivated virus. Cell viability was quantified by measuring absorbance at 450 nm on a VersaMax microplate reader (Molecular Devices).

### Caspase-1 and Caspase-3 Assays

10^6^ A549 cells were infected with ZIKV-WT, TL.PK, and p.2.5’ viruses at an MOI of 0.5 for one hour at 37°C. After infection, inoculum was removed, cells were washed three times with 1 mL 1X PBS, and 2 mL of complete Ham’s F-12 media. Samples were harvested at 24-, and 48-hours post-infection. Caspase activation was assayed with the Caspase-3 and Caspase-1 colorimetric assay kits from abcam (ab39401 and ab273268, respectively). At harvest, culture media was removed, and cells were washed one with 1X PBS. Cells were then scraped in 2 mL of 1 mL of 1X PBS and pelleted at 2500 x g for five minutes at 4°C. Cell pellets were washed twice with 1 mL 1X PBS and pelleted again as previously stated. After the final wash, pellets were resuspended in 50 μL of Cell Lysis Buffer and incubated on ice for 10 minutes. Cell debris were pelleted at 10,000 x g for one minute at 4°C. Lysate samples were flash frozen and stored at −80°C until time of assay. Lysate protein concentrations were quantified using the Pierce™ BCA Protein Assay Kit from ThermoFisher. Activated caspase-3 and caspase-1 were assayed following the manufacturer’s protocols from each caspase assay kit used.

### Northern blot

A549 cells were plated at a density of 2e^5^ cells per well in 6-well cell culture plates. Plates were infected with ZIKV-WT, TL.PK, and p.2.5’ viruses at an MOI of 0.5. Cells were infected with 500 μL of viral inoculum for one hour at 37°C. After infection, inoculum was removed, cells were washed three times with 1 mL 1X PBS, and 2 mL of complete Ham’s F-12 media. Samples were harvested at 0-, 24-, and 48-hours post-infection. At harvest, media was aspirated, and cells were washed three times with 1X PBS. Three wells were combined for each strain at each time point to obtain sufficient levels of RNA. Total RNA was extracted from infected cells using the OmegaBioTek E.Z.N.A Total RNA Kit I. RNA was extracted according to the manufacturer’s protocol. Northern blot for ZIKV sfRNA was performed according to Akiyama 2021^17^. Densitometry analysis was performed according to previously published protocols^37^.

### In Vitro Xrn-1 Digest and Analysis

Methods adapted from Slonchack 2022^34^. ZIKV 3’UTR templates were PCR amplified from the p2 plasmid using primers that includes 100-nt of NS5 as an Xrn-1 leader sequence, plus a T7 RNA promoter sequence to the 5’ end of the template (Table 3). A control RNA was amplified from the NS5 gene upstream of the 3’UTR-amplyfiying forward primer (Table 3). RNA was in vitro transcribed using the NEB HiScribe T7 Arca kit, incubating the reaction for 1 hour. Following in vitro transcription RNA was purified using the Zymo RNA Clean and Concentrator kit, eluting in 25 μL. 2 μg of RNA was folded in NEBuffer3 at 90°C for 3 minutes, followed by 20°C for 5 minutes. In vitro Xrn-1 digests were performed with 1U Xrn-1 (NEB), 10U RppH (NEB), and 1U RNAseOUT (Invitrogen). Half of the RNA folding reaction was used in an Xrn-1 positive digest, while the other half was used for an Xrn-1 negative digest. Digests were incubated at 37°C for 2 hours.

**Table 3.**
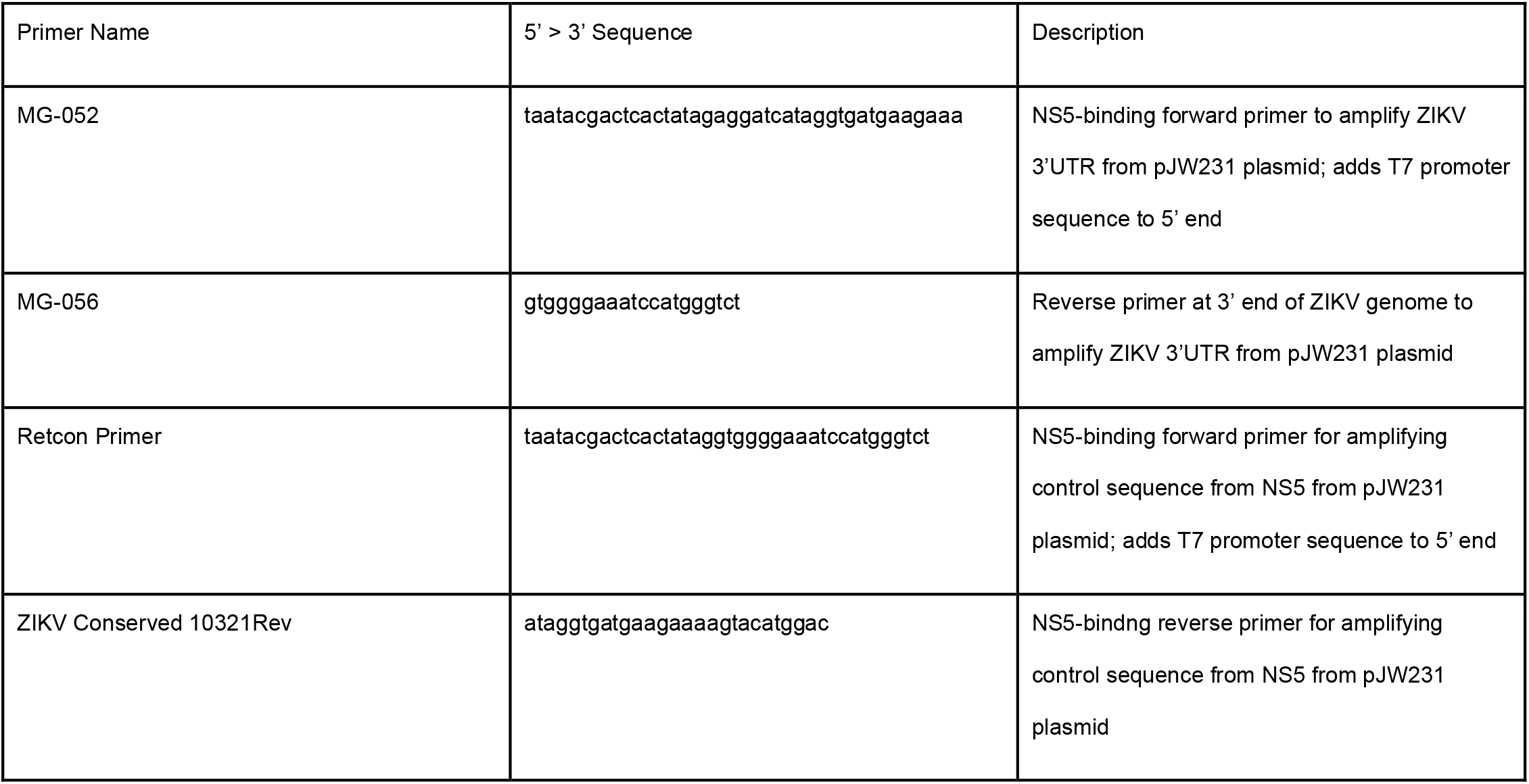
XRN1 Digest Assay Primers.

Digest products were analyzed by gel electrophoresis. A 5% TBE-Urea gel (BioRad) was pre-run at 200 V for 15 minutes in 1X TBE buffer. Xrn-1 digest samples were diluted 1:1 in 2X RNA loading dye (BioRad) and RNA was denatured at 80°C for 90 seconds. Samples were run on the gel at 150V for 45 minutes in 1X TBE. The gel was stained using 0.5 μg/mL ethidium bromide in 1X TBE for 30 minutes. Gels were imaged with a G:Box gel imager (Syngene).

### ISG RNA Expression

A549 cells were infected at an MOI = 1 as described above. For poly I:C controls, 10^6^ were treated with 50 μg/mL of poly I:C in complete Ham’s F-12 media (described above). At 24 hours post-infection or post-treatment, RNA was extracted from cells using the OmegaBiotek E.Z.N.A Total RNA Extraction Kit following manufacture’s protocol. Whole cell RNA was reverse transcribed using the BioRad gDNA Clear cDNA Synthesis kit with 100 ng of RNA template and the addition of 1 μL of (RT control from plate) to each reaction. qPCR was done with BioRad PrimePCR Interferon α/β Signaling Pathway plates according to manufacturer’s protocol.

### Interferon Dose Response Curve

10^4^ Vero cells were infected with ZIKV-WT, -TL.PK, -P.2.5’, or mock infected at an MOI=1 for one hour at 37°C. After infection, cells were treated with IFN-β with a 1:2 dilution series from 100 – 3.125 IU/mL, including cells that went untreated. After 48 hours supernatant was harvested for viral RNA extraction. Viral RNA was extracted using the Omega-Biotek E.Z.N.A Viral RNA extraction kit. Viral genome copies in the supernatant were quantified using RT-qPCR as described above. interferon-β treatments were done in triplicate for each virus condition.

### Animal Studies

All animal and infectious disease studies were reviewed and approved by the University of Colorado Institutional Animal Care and Use Committee and Institutional Biosafety Committee. B6.CG-*Ifngr1^tm1Agt^Ifnar1^tm1.2Ees^/J* mice purchased from Jackson Laboratories (JAX stock #029098) were maintained and bred in specific-pathogen-free facilities at the University of Colorado Anschutz Medical Campus animal facility. Animals to be infected were housed in an animal BSL-3 (ABSL-3) facility. After infection, mice were observed daily for weight loss and signs of disease until the experiment endpoint.

### Murine Experiments

6–8-week-old mice were anesthetized with isoflurane and infected via intraperitoneal injection with 10^4^ PFU of virus. For mice assayed for survival curve, infected individuals were weighed daily until they reached 80% of their original weight at time of infection. Mice were also monitored for other symptoms of illness, including hunched body, slow movement, and matted fur. Once mice showed significant illness and reached the weight cutoff they were euthanized with isoflurane.

Animals infected for measuring viral load were euthanized with isoflurane 4-, 6-, or 8-days post-infection. Mice were perfused with 25 – 30 mL of 1X PBS, followed by dissection and harvest of spleen and brain. Organs were preserved in RNALater solution until RNA extraction. RNA was extracted from tissue following the techniques used in Sparks 2020. 1 μg of total organ RNA was reverse transcribed using the BioRad iScript reverse transcription kit according to manufacturer’s protocol. qPCR for Zika genome copies was performed as described above. Viral load was normalized by dividing the qPCR starting quantity value by the mass of harvested organ for each sample.

### Statistical Analysis

Data was analyzed by two-way ANOVA, assuming normal distribution. AG129 survival curve was analyzed using Kaplan-Meyer graph. All statistical analysis done using GraphPad Prism.

## Notes

### Competing Interest Statement

The authors have declared no competing interest.

